# The transcription factor *RXRA* regulates lipid metabolism in duck myoblasts by the *CD36* network

**DOI:** 10.1101/2022.04.13.488167

**Authors:** Ziyi Pan, Guoyu Li, Guoqing Du, Dongsheng Wu, Xuewen Li, Yu Wang, Junxian Zhao, Xiran Zhang, Xingyong Chen, Chen Zhang, Sihua Jing, Zhaoyu Geng

## Abstract

Lipid metabolism is a highly complex metabolic process regulated at multiple levels. RXRA is a well-characterized factor that regulates lipid metabolism. To explore the function and mechanism of the transcription factor *RXRA* in myoblasts, and to further explore the key factors that *RXRA* regulates the target gene *CD36* signaling network to regulate lipid metabolism. We found that the transcription factor *RXRA* inhibited the accumulation of triglycerides (TGs), cholesterol (CHOL) and non-esterified fatty acids (NEFA) in CS2 cells by regulating CD36 as well as the fatty acid beta oxidation pathway. CD36 functions similar to *RXRA* in myoblasts. *CD36* overexpression reduced the levels of TGs, CHOL, NEFAs and saturated fatty acids (SFAs) in these cells, while *CD36* knockout increased the levels of TGs, CHOL, NEFAs, SFAs and monounsaturated fatty acids (MUPAs) in these cells. *GRB2*, *MAP1B*, *SLC38A1*, *SLC4A7*, *NCOA3*, *PKIA*, *MOB2*, *SAA2* and *RXRA* are involved in the *CD36* promotion of lipid efflux through lipid metabolism, endocytosis and amino acid metabolism. Considering these results, we propose that *RXRA* regulates lipid metabolism in myoblasts by regulating the *CD36* signaling network.

## Introduction

In the past 20 years, the world’s demand for poultry meat protein has continued to increase, and the need for duck meat has also increased. According to reports, Asia produced nearly 3 million tons of duck meat in 2017, accounting for 75% of the world’s production (Gariglio *et al*, 2021). The quality of duck meat is an essential factor influencing consumer preferences. Intramuscular fat (IMF) content is an important factor in determining duck meat quality because it affects duck meat flavour, juiciness, tenderness, and muscle colour (Fan *et al*, 2020). Duck IMF is primarily composed of triglycerides (TGs) and cholesterol (CHOL) (Daniel *et al*, 2018), and fatty acids (FAs) play major roles in the nutritional and product quality of meat(Wood & Enser, 2017). Increasing the contents of unsaturated FAs in the human diet can reduce the risk of cardiovascular disease and diabetes (Gillingham *et al*, 2011; Laaksonen & David, 2005). However, too many FAs are key factors leading to obesity and related diseases. The excessive accumulation of FAs in the body is known to reduce glucose utilization and decrease glucose uptake (Randle *et al*, 1963). The collection of FAs in skeletal muscle can cause a mitochondrial emergency in cells and insulin resistance, leading to overall cell dysfunction(Tumova *et al*, 2015). Therefore, controlling the FA metabolism of meat ducks has become an important research topic in myoblast fat accumulation and human health diet in recent years.

Transcription factors are among the molecules regulating lipid metabolism. Retinoid X receptors (RXRs) and peroxide proliferator-activated receptors (PPARs) are ligand-activated transcription factors that coordinate and regulate gene expression. This PPAR-RXR transcription complex plays a key role in TG metabolism, FA removal, and glucose homeostasis (Plutzky & J., 2011). Heterozygous PPARG-deficient mice were treated with RXR antagonists and lost white fat tissue and presented with significantly reduced leptin and energy expenditures, which led to increases in TG content in skeletal muscle and insulin resistance (Yamauchi *et al*, 2001). It has been reported that RXR agonists are insulin sensitizers and can reduce blood sugar, hypertriglyceridaemia and hyperinsulinaemia (Mukherjee *et al*, 1997). The reduced fat gain in RXRG-knockout mice fed a high-fat diet indicated that RXR plays an important role in the common diseases of dyslipidaemia and obesity (Haugen *et al*, 2004). However, the effect and mechanism of *RXRA* on lipid metabolism in duck myoblasts have not been extensively studied. Therefore, the functions of *RXRA* and the related molecular mechanisms through which they regulate target gene expression must be systematically studied.

*CD36* is a multifunctional membrane receptor involved in FA uptake, lipid metabolism, atherosclerosis and inflammation (Ulug & Nergiz-Unal, 2020). Acute overexpression of *CD36* in human skeletal muscle cells significantly increased downstream insulin-stimulated Akt phosphorylation and FA uptake, the oxidation rate, and total lipid accumulation (Bell *et al*, 2010). Compared with wild-type mice, *CD36* transgenic mice showed reduced muscle fat, blood TG and FA contents (Ibrahimi *et al*, 1999). Depalmitoylated CD36 recruits a different tyrosine kinase, SYK, which phosphorylates VAV and JNK to internalize pits and transport FAs to the endoplasmic reticulum for esterification and transport to adipocytes for storage(Hao *et al*, 2020). Therefore, exploration of factors that affect *CD36* expression is crucial to addressing lipid metabolism disorders in myoblasts. Previous data have shown that *CD36* is a differentially expressed gene (DEG) in ducks with low and high residual feed intake (RFI) and that RFI is affected by lipid metabolism (Rauw *et al*, 2012; Richardson *et al*, 2004). A prediction study of the transcription factor promoter binding region suggested that the promoter region of *CD36* has five *RXRA*-binding sites.

Based on these findings, multiple sets of analytical methods were used in this study to explore *RXRA*, and the results revealed that *RXRA* is involved in the lipid metabolism of myoblasts. *RXRA* is a transcription factor of *CD36*, which also regulates gene expression. Further experiments showed that *CD36* exhibited functions similar to those of *RXRA*; for example, they both regulate lipid metabolism in myoblasts and maintain cell stability through FA metabolism, amino acid metabolism and the endocytic pathway. In conclusion, our study systematically expounds the regulatory network of *RXRA* that affects the lipid metabolism of duck myoblasts and suggests new strategies for approaching the issue of myoblast fat accumulation and human health diet.

## Results

### Trends in RXRA expression

Signaling factors play an indispensable role in regulating lipid metabolism in myoblasts(Liu *et al*, 2020). *RXRA* is a key transcriptional regulator in the regulation of intramuscular fat metabolism in livestock and poultry. *RXRA* has distinct expression patterns during the stages of intramuscular adipocyte differentiation(Zhang *et al*, 2017). In order to explore the function and mechanism of transcription factor *RXRA* in duck myoblasts, we highly expressed and knocked down the duck *RXRA* gene in myoblasts. After successful construction, the oeRXRA, oeNC, shRXRA, and shNC vectors were transferred to the cell culture plate when the CS2 cells reached 80% confluency. After 24 h of transfection, the cell morphology and the expression of green fluorescence protein were observed with an inverted fluorescence microscope (Fig 1A). The cells were collected, and RNA was extracted. The expression of *RXRA* mRNA in CS2 cells was detected with RT–PCR. The results showed that, compared with cells transfected with an empty vector, *RXRA* was effectively overexpressed and knocked down after oeRXRA and shRXRA transfection, respectively (Fig 1B). Then, western blot analysis was performed to verify the level of RXRA protein expression after the transfection of oeRXRA and shRXRA at 24 h. RXRA overexpression in CS2 cells transfected with oeRXRA was confirmed by western blot analysis (Fig 1C; Appendix Fig S1). Moreover, RXRA expression was effectively knocked down after shRXRA transfection (Fig 1C; Appendix Fig S1). Sequence alignment of the *RXRA* overexpression and interference vectors is shown in Appendix Fig S2. The results of the screening of the four interference vectors against *RXRA* are shown in Appendix Fig S3.

**Figure 1.**
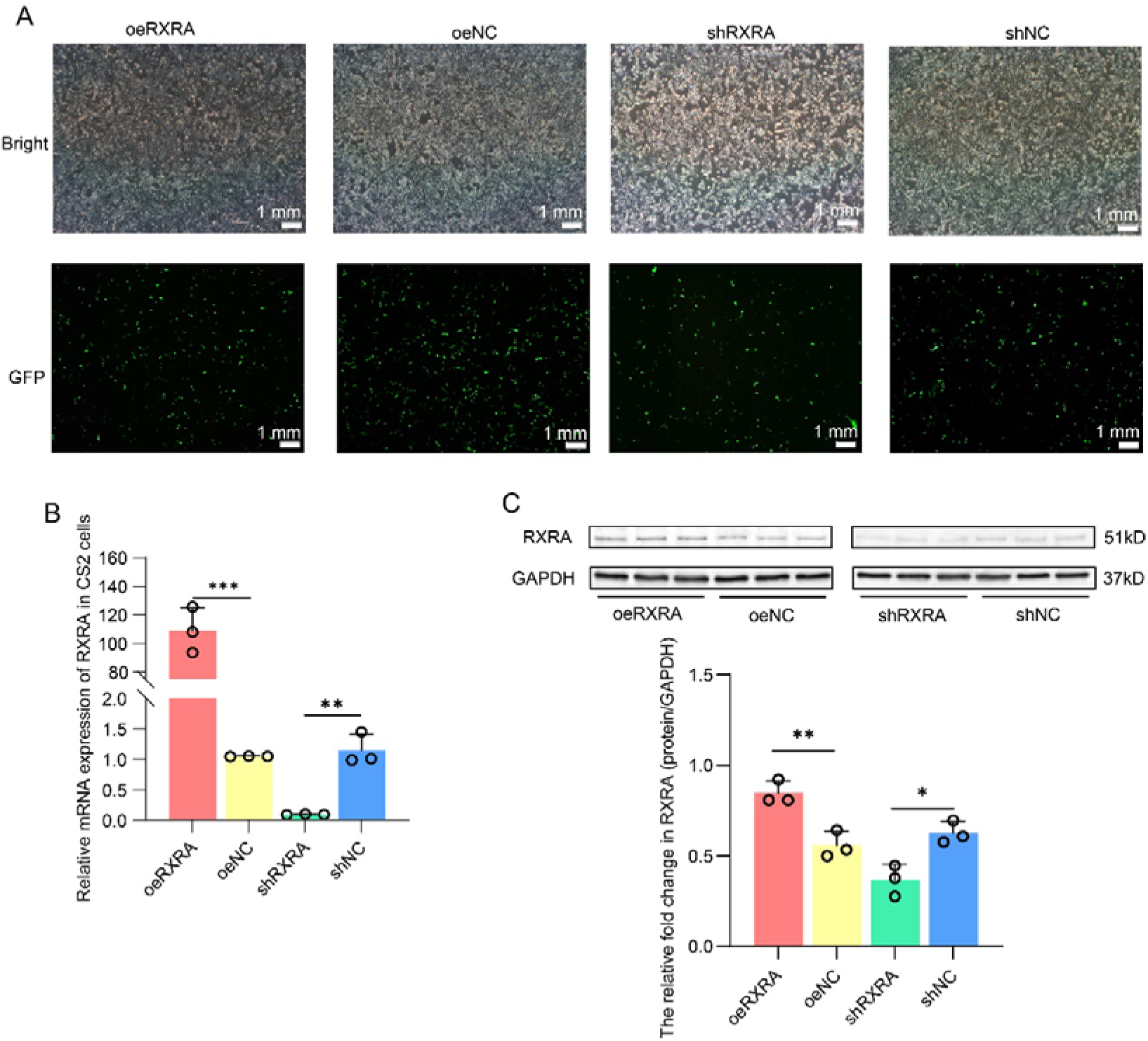
The mRNA and protein expression levels of *RXRA* in CS2 cells after transfection with oeRXRA, oeNC, shRXRA and shNC. A. Microscopy observation of CS2 cells transfected with oeRXRA, oeNC, shRXRA, and shNC. B. Relative mRNA expression levels of *RXRA* in CS2 cells transfected with oeRXRA, oeNC, shRXRA, and shNC (data are shown as the means ± SD, n=3 independent experiments, ** *P* < 0.01, *** *P* < 0.001, t-test). C. The indicated protein levels were detected by western blot analysis. The relative fold change in RXRA (protein/GAPDH) as determined by western blotting was quantified through a greyscale scan (data are shown as the means ± SD, n=3 independent experiments, \**P* < 0.05, ** *P* < 0.01, t-test).

### The TG, CHOL, and NEFA contents

After the *RXRA* gene was knocked out in mice, serum triglyceride and cholesterol levels were reduced(RaŸny *et al*, 2010). The diversity of organisms leads to the fact that the specific functions of genes cannot be generalized. Different organizations and modes of action produce different results(Mukherjee *et al*, 1997; Haugen *et al*, 2004). The overexpression vectors oeRXRA and oeNC and the short hairpin (interference) vectors shRXRA and shNC were transfected into CS2 cells. The cells were collected 24 h later, and the TG, CHOL, and NEFA contents in the CS2 cells were detected according to the respective kit instructions. Compared with that in the normal control vector group, the TG content in the oeRXRA group was significantly reduced, and the TG content in the shRXRA group was significantly increased (Fig. 2A). *RXRA* has been previously shown to play a key role in CHOL homeostasis, intestinal CHOL absorption, and bile acid synthesis(Willy *et al*, 1995). The CHOL content in the oeRXRA group was significantly lower than that in the oeNC group, while the CHOL content in the shRXRA group did not change significantly (Fig. 2B). In addition, we detected changes in the NEFA content in CS2 cells, and the results showed that the NEFA content in the oeRXRA group was significantly reduced compared with that in the oeNC group. In contrast, the NEFA content in the shRXRA group was significantly higher than that in the shNC empty vector group (Fig. 2C).

**Figure 2.**
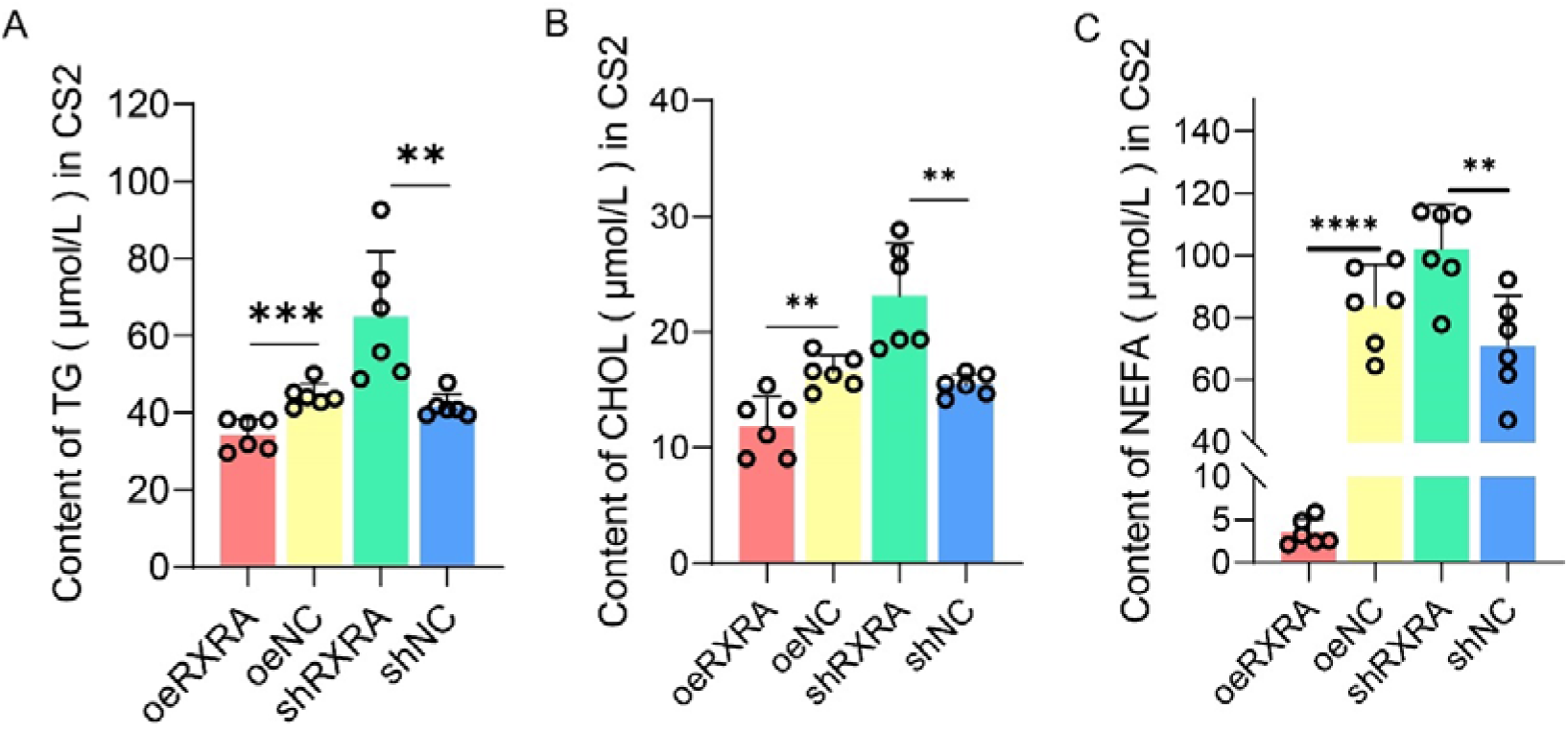
Effect of *RXRA* on the triglyceride (TG), cholesterol (CHOL), and nonesterified fatty acid (NEFA) contents in CS2 cells. A. TG contents in CS2 cells transfected with oeRXRA, oeNC, shRXRA, and shNC (data are shown as the means ± SD, n=6 independent experiments, ** *P* < 0.01, *** *P* < 0.001, t-test). B. CHOL contents in CS2 cells transfected with oeRXRA, oeNC, shRXRA, and shNC (data are shown as the means ± SD, n=6 independent experiments, ** *P* < 0.01, t-test). C. NEFA contents in CS2 cells transfected with oeRXRA, oeNC, shRXRA, and shNC (data are shown as the means ± SD, n=6 independent experiments, ** *P* < 0.01, **** *P* < 0.0001, t-test).

### Relative expression levels of lipid metabolism genes

RT–PCR was performed to detect the expression of lipid metabolism-related genes after *RXRA* interference and overexpression. As shown in Figure 3 A, the relative expression levels of the *CD36*, *ACSL1* and *PPARG* genes in the oeRXRA group were significantly higher than those in the oeNC group, and there was no change in the expression levels of the *FABPA* or *ELOVL6* gene between the two groups. The *CD36*, *ACSL1*, *ELOVL6* and *PPARG* gene expression levels in the shRXRA group were significantly higher than those in the shNC group, and there was no significant change in the *FABPA* gene expression level between the two groups (Fig. 3A). The results of the western blot analysis were consistent with the results of the gene mRNA expression analysis (Fig. 3B; Appendix Fig S4). *RXRA* usually functions in the form of RXR/PPAR complex in organisms(Deng *et al*, 2019). *PPARG* can induce the expression of *ACSL 1*, which plays an important role in FA breakdown(Bervejillo *et al*, 2020). RXRA may promote the activation of fatty acid beta oxidation pathway by binding to PPARG in myoblasts. We found that when we changed the expression of the *RXRA* gene, the expression of *CD36* changed to the greatest degree among the genes analysed. Therefore, we speculated that there may be a regulatory relationship between *RXRA* and *CD36*. We used PROMO software to predict five *RXRA*-binding sites in the 2000-bp promoter region of the target gene *CD36*, which is expressed downstream of the *RXRA* transcription factor (Fig. 3C). The results of cotransfection showed that oeRXRA increased the wt-CD36 promoter luciferase activity compared with transfection of mu-CD36, GLP3-Basic or oeNC. However, transfection of oeNC exerted no significant effect on wt-CD36, mu-CD36 or GLP3-Basic luciferase activities (Fig. 3D). The overexpression of RXRA increased CD36 promoter activity through five targets: TGACCTCAT, CAAAGGTCA, CAGAGGTCA, TGACCTGTT and TGACCTTGG. Sequence alignment of wt-CD36 and mu-CD36 vectors is shown in Appendix Fig S5.

**Figure 3.**
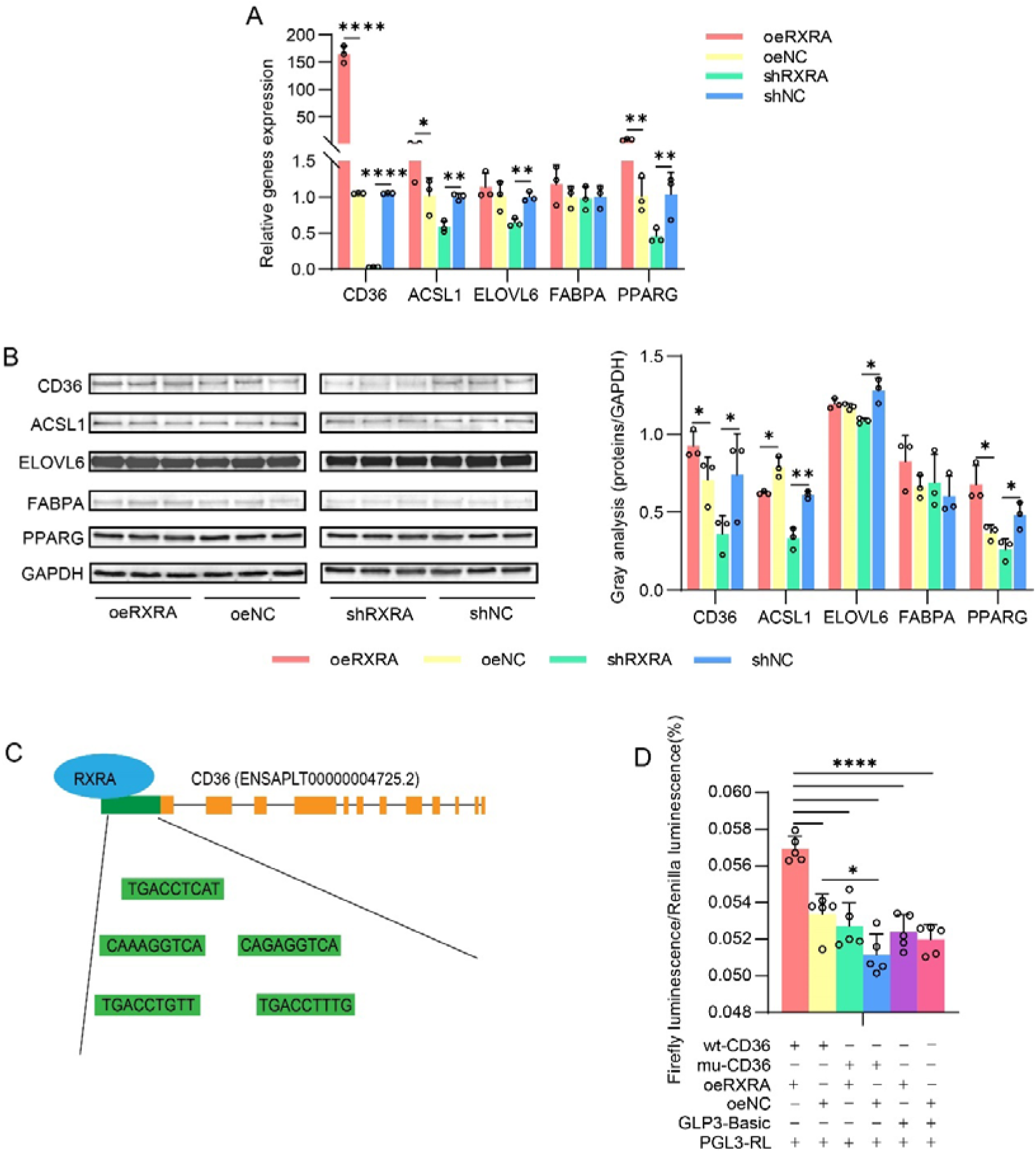
Effect of *RXRA* on lipid metabolism-related gene and protein expression in CS2 cells. A. The gene expression levels of *CD36*, *ACSL1*, *ELOVL6*, *FABPA* and *PPARG* after transfection of overexpression vectors oeRXRA, oeNC, shRXRA and shNC in CS2 cells (data are shown as the means ± SD, n=3 independent experiments, \**P* < 0.05, ** *P* < 0.01, and **** *P* < 0.0001, t-test). B. The indicated protein levels were detected by western blotting. The relative changes in CD36, ACSL1, ELOVL6, FABPA, and PPARG expression (protein/GAPDH) in the western blots were quantified by the greyscale scan and reported as the fold change (data are shown as the means ± SD, n=3 independent experiments, \**P* < 0.05, ** *P* < 0.01, t-test). C. Prediction of RXRA-binding sites in the *CD36* promoter region. D. Analysis of firefly luciferase activity and Renilla luciferase activity (data are shown as the means ± SD, n=5 independent experiments, **** *P* < 0.0001, t-test).

### Trends in CD36 expression

The above research results found that *RXRA* can bind to the *CD36* promoter sequence to promote its expression. *CD36* plays an important role in fatty acid transport in adipocytes, which provides an important downstream gene for *RXRA* to precisely regulate lipid metabolism in myoblasts(Hao *et al*, 2020). Moreover, *CD36*, which function as a fatty acid transporter, lipoprotein receptor, and signaling molecule (Su & Abumrad, 2009), plays a key role in this process (Bartelt *et al*, 2011). To explore whether *CD36* is related to *RXRA* gene function, we overexpressed and interfered with *CD36* in myoblasts. After successful construction, the oeCD36, oeNC, shCD36, and shNC vectors were transferred to the cell culture plate when the CS2 cells reached 80% confluency. After 24 h of transfection, the cell morphology and the expression of the green fluorescence protein were observed under an inverted fluorescence microscope (Fig. 4 A). The cells were collected, and RNA was extracted. The expression of *CD36* mRNA in CS2 cells was detected by RT–PCR. The results showed that compared with cells transfected with the empty vector, *CD36* expression was effectively overexpressed and knocked down after oeCD36 and shCD36 vector transfection (Fig. 4 B). Then, western blot analysis was performed to verify the level of CD36 protein expression 24 h after the transfection of oeCD36 and shCD36. CD36 overexpression was confirmed in CS2 cells transfected with oeCD36 by western blot analysis (Fig. 4C; Appendix Fig S6). In addition, CD36 expression was effectively knocked down after shCD36 vector transfection. (Fig. 4C; Appendix Fig S6). The sequence alignment of *CD36* overexpression and interference vectors is shown in Appendix Fig S7. The results from screening the four interference vectors against *CD36* are shown in Appendix Fig S8.

**Figure 4.**
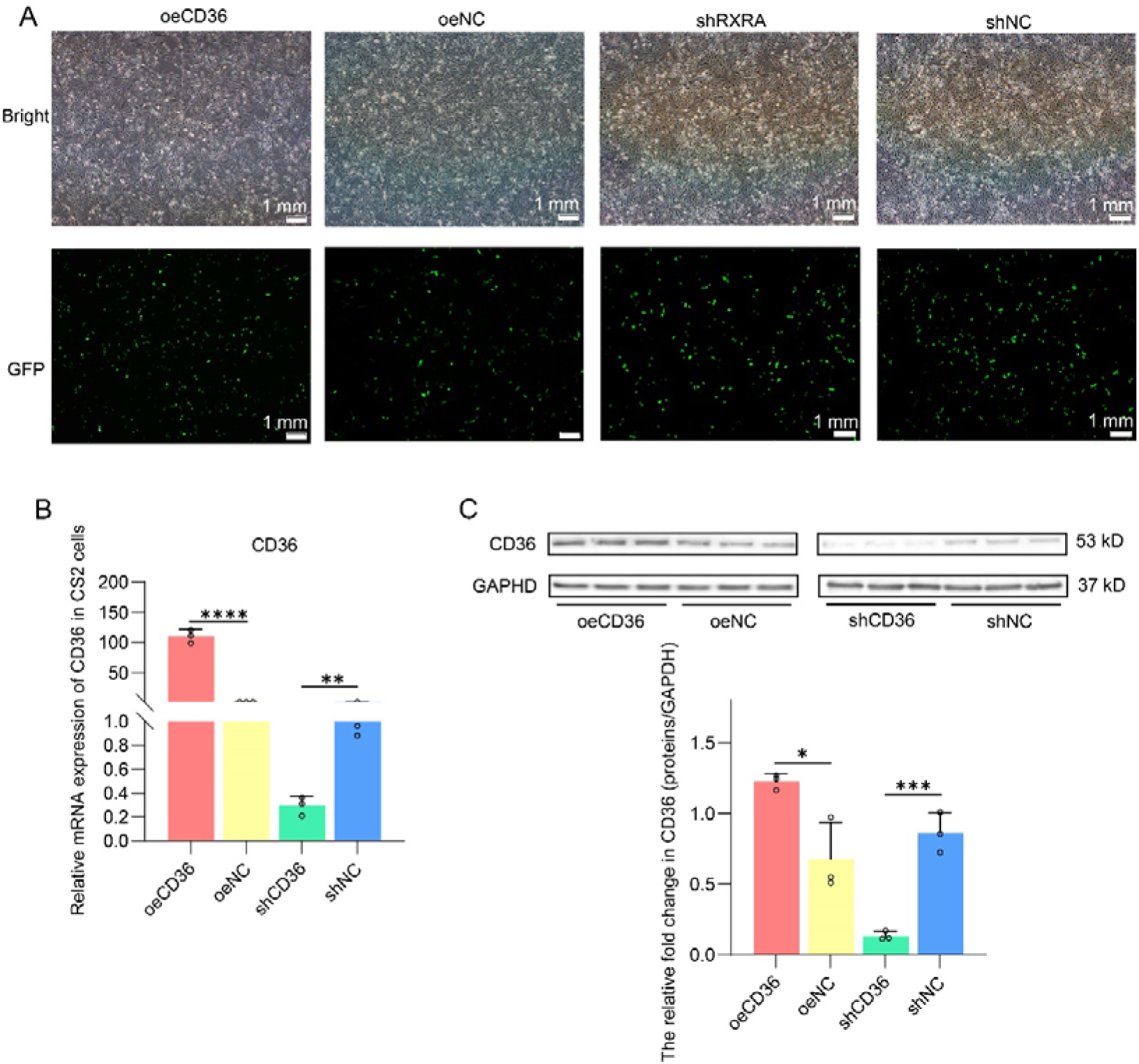
The mRNA and protein expression levels of *CD36* in CS2 cells after transfection with overexpression vectors oeCD36, oeNC, shCD36 and shNC. A. Microscopy observation of CS2 cells transfected with oeCD36, oeNC, shCD36, and shNC. B. Relative mRNA expression levels of *CD36* in CS2 cells transfected with oeCD36, oeNC, shCD36, and shNC (data are shown as the means ± SD, n=3 independent experiments, ** *P* < 0.01, **** *P* < 0.0001, t-test). C. The indicated protein levels were detected by western blotting. The relative fold change in CD36 (protein/GAPDH) expression as detected by western blotting was quantified through a greyscale scan (data are shown as the means ± SD, n=3 independent experiments, \**P* < 0.05, *** *P* < 0.001, t-test).

### CD36 alters the lipid composition in CS2 cells

Transfection of oeCD36, oeNC, shCD36, and shNC into CS2 cells. The cells were collected 24 h later, and the TG, CHOL, and NEFA contents in the CS2 cells were measured according to the respective kit instructions. *CD36* and *RXRA* play the same roles in CS2 cells. Compared with the overexpression and interference empty vector control groups, the TG content in the oeCD36 group was significantly reduced, and the TG content in the shCD36 group was significantly increased (Fig. 5A). *CD36* reduced lipid accumulation in HepG2 cells in a nonalcoholic fatty liver treatment strategy similar to our findings(Shen *et al*, 2021). The CHOL content in the oeCD36 group was significantly lower than that in the oeNC group, while the CHOL content in the shRXRA group was significantly increased (Fig. 5B). After high-fat intake, the level of cholesterol in the liver of mice increased, accompanied by an increase in the expression of *CD36*, which plays an important role in maintaining cholesterol balance in the body (Nergiz-Unal *et al*, 2020). *CD36* is able to take up FAs through a dynamic palmitoyl-regulated endocytic pathway(Hao *et al*, 2020). We analysed changes in the NEFA content in CS2 cells, and the results showed that the NEFA content in the oeCD36 group was significantly lower than that in the oeNC group. In contrast, the NEFA content in the shCD36 group was significantly higher than that in the shNC group (Fig. 5C). Notably, long-chain FAs are essential for lipid metabolism(Tran *et al*, 2016). To further explore the mechanism by which *CD36* regulates lipid metabolism in CS2 cells, an FA methyl ester mixture containing 38 components was measured via gas chromatography and mass spectrometry (GC–MS). The overexpression of the *CD36* gene in CS2 cells led to a decrease in the SFA content, while the knockdown of the *CD36* gene increased the SFA content (Fig. 5D). The contents of C12:0, C14:0 and C18:0 in the oeCD36 group were significantly lower than those in the oeNC group, and the contents of C12:0, C14:0, C16:0 and C18:0 in the shCD36 group were significantly higher than those in the shNC group (Fig. 5E). The overexpression of the CD36 gene in CS2 cells did not change the MUFA content, but knocking down the expression level of the CD36 gene promoted the accumulation of MUFAs (Fig. 5D). Notably, oeCD36 promoted greater excretion of C15:1, C17:1 and C18:1N9C than oeNC, but shCD36 inhibited the excretion of C16:1, C18:1N12T, C18:1N12 and C18:1N9C compared with shNC (Fig. 5F). Finally, the expression of CD36 in CS2s did not change the content of PUFAs (Fig. 5D), but shCD36 promoted an increase in the contents of C18:2N6, C20:4N6, and C22:5N3 compared with those in the shNC group (Fig. 5G).

**Figure 5.**
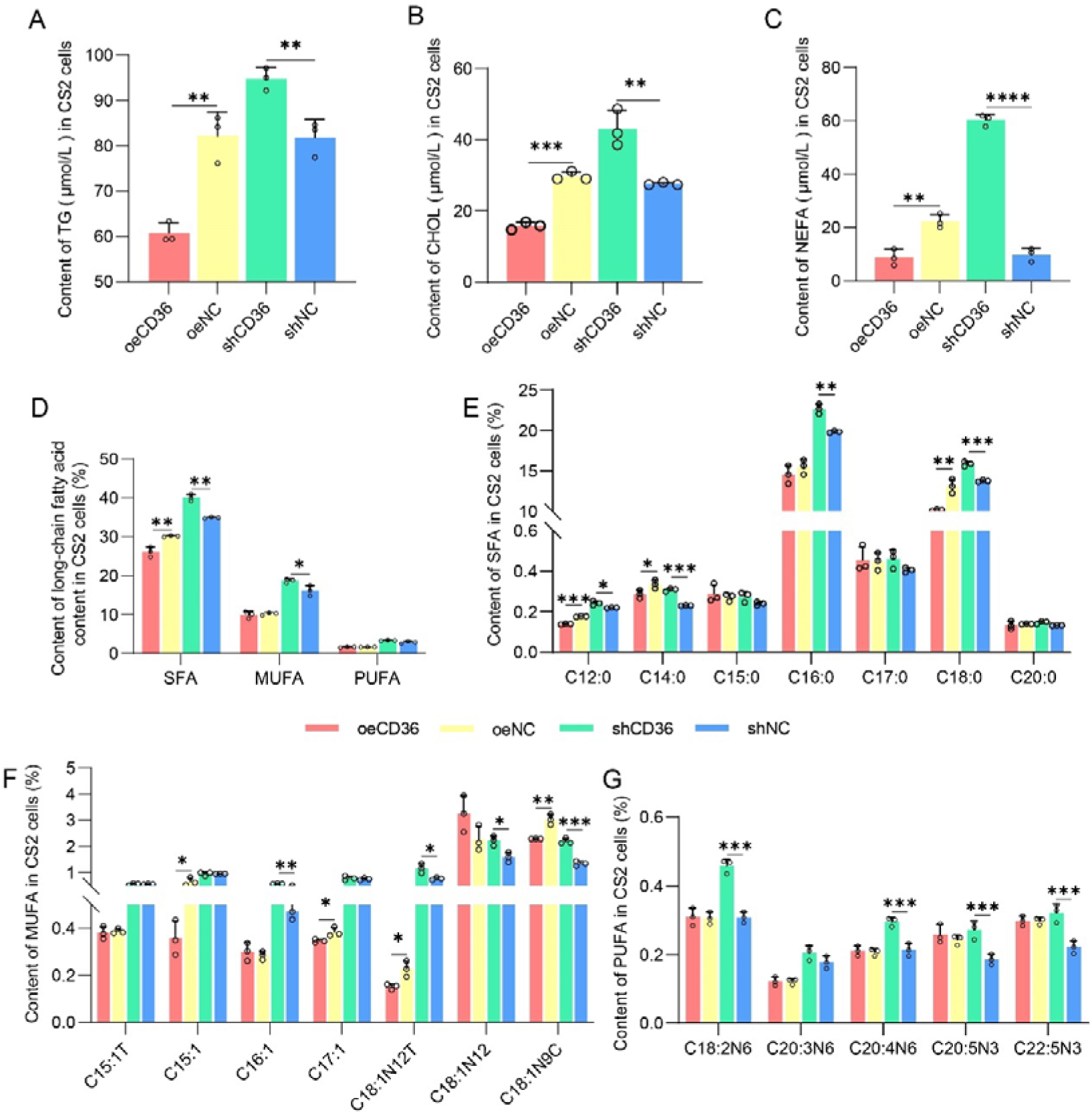
Detection of the triglyceride (TG), cholesterol (CHOL), and nonesterified fatty acid (NEFA) contents and fatty acid (FA) composition in CS2 cells. A. TG content in CS2 cells after transfection with oeCD36, oeNC, shCD36 or shNC (data are shown as the means ± SD, n=3 independent experiments, ** *P* < 0.01, t-test); B. CHOL content in CS2 cells after transfection with oeCD36, oeNC, shCD36 or shNC (data are shown as the means ± SD, n=3 independent experiments, ** *P* < 0.01, *** *P* < 0.001, t-test); C. NEFA content in CS2 cells after transfection with oeCD36, oeNC, shCD36 or shNC (data are shown as the means ± SD, n=3 independent experiments, ** *P* < 0.01, **** *P* < 0.0001, t-test). D. Fatty acid (FA) composition in CS2 cells after transfection with oeCD36, oeNC, shCD36 or shNC (data are shown as the means ± SD, n=3 independent experiments, \**P* <0.05, ** *P* < 0.01, t-test). E. The saturated fatty acid (SFA) content after transfection of oeCD36, oeNC, shCD36 and shNC (data are shown as the means ± SD, n=3 independent experiments, \**P* <0.05, ** *P* < 0.01, *** *P* < 0.001, t-test). F. The monounsaturated fatty acid (MUFA) content after transfection of oeCD36, oeNC, shCD36 and shNC (data are shown as the means ± SD, n=3 independent experiments, \**P* <0.05, ** *P* < 0.01, *** *P* < 0.001, t-test). G. The polyunsaturated fatty acid (PUFA) content after transfection of oeCD36, oeNC, shCD36 and shNC (data are shown as the means ± SD, n=3 independent experiments, *** *P* < 0.001, t-test).

### DEGs in CS2 cells with CD36 overexpression and knockdown

From the above results, we can find that *RXRA* can reduce lipid accumulation in myoblasts by precisely regulating the expression of *CD36*. To deeply mine the regulatory network downstream of the *RXRA* target gene *CD36*, RNA-seq is an important tool for this purpose. RNA-seq data were obtained from samples of CS2 cells 24 h after transfection with oeCD36 or shCD36 plasmids. A PCA of the transcriptome sequencing data obtained from 6 CS2 cell samples (Fig. S9) revealed that the gene expression in two groups of these cells clustered together. A total of 120 DEGs were identified in CS2 cells transfected with oeCD36 or shCD36. Compared with the shCD36 group, the oeCD36-transfected group had 23 upregulated and 97 downregulated genes (Fig. 6; Table EV1). The set of DEGs between the oeCD36 and shCD36 groups included a large number of genes related to lipid transport and lipid metabolism (Table 1).

**Figure 6.**
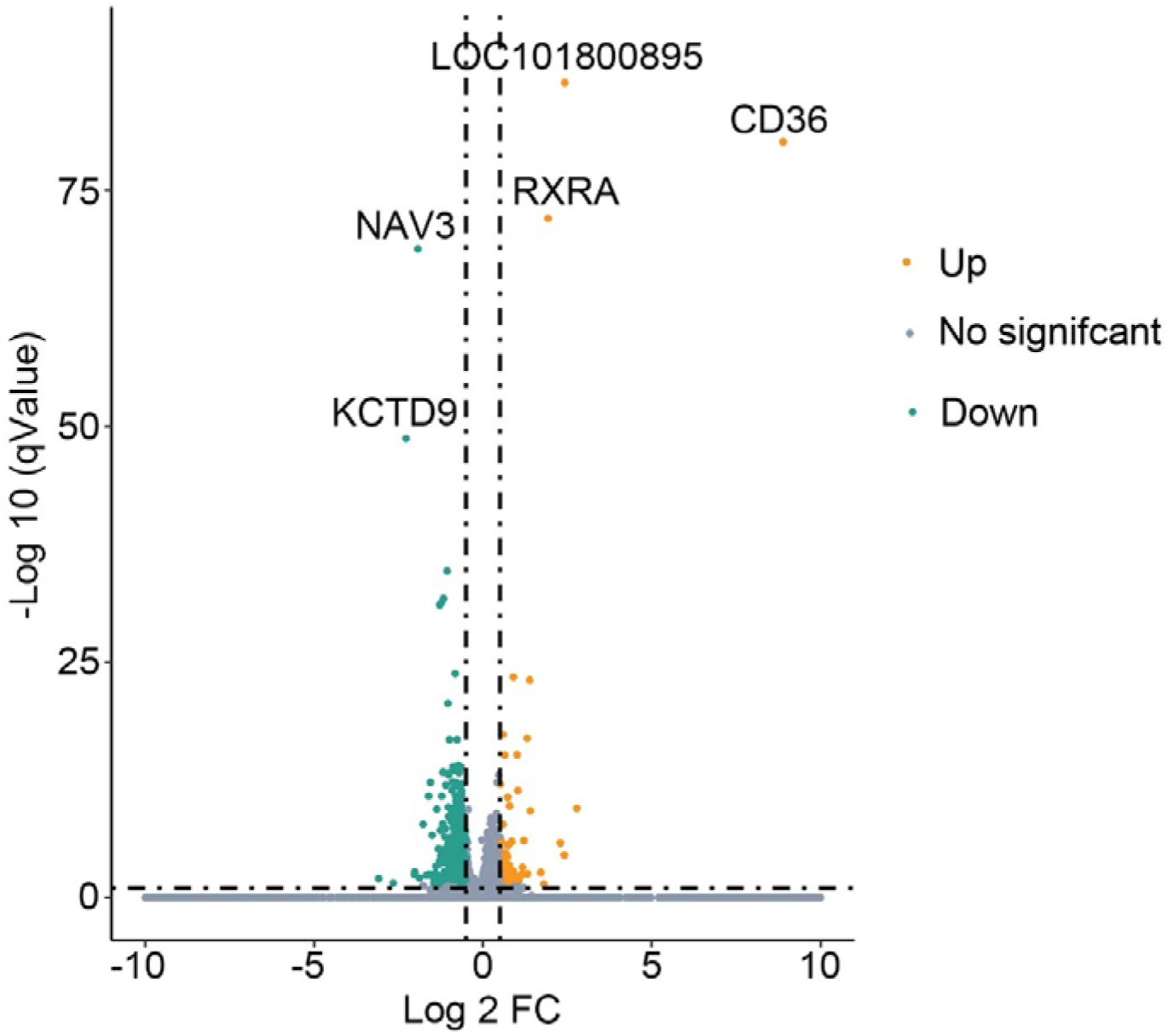
Volcano plot showing differentially expressed genes (DEGs). The scale and significance of the DEGs between the oeCD36 and shCD36 groups are shown. Orange dots represent upregulated genes, and cyan dots represent downregulated genes (the criteria for DEG designation were Log2 FC > 1 or Log2 FC < -1, Q-value < 0.05).

**Table 1.** A part of DEGs between oeCD36 and shCD36 CS2 cell related to lipidmetabolism

| Gene ID | Log2FC | Q-value | oeCD36 | shCD36 | Trend |
| --- | --- | --- | --- | --- | --- |
| GRB2 | -1.29 | 0.04 | 45.12 | 110.91 | Down |
| MAP1B | -1.28 | 9.14E-32 | 10.74 | 26.02 | Down |
| SLC38A1 | -1.11 | 0.01 | 57.41 | 123.59 | Down |
| SLC4A7 | -1.07 | 0.01 | 5.21 | 10.92 | Down |
| NCOA3 | -1.03 | 4.17E-07 | 5.95 | 12.17 | Down |
| PKIA | 1.04 | 0.02 | 21.58 | 10.47 | Up |
| MOB2 | 1.08 | 0.01 | 131.1 | 62.22 | Up |
| SAA2 | 1.31 | 1.22E-17 | 127.8 | 51.46 | Up |
| CD36 | 8.89 | 6.22E-81 | 25.9 | 0.05 | Up |
| RXRA | 1.93 | 8.79E-73 | 22.11 | 5.82 | Up |

### GO and KEGG enrichment analyses of the DEGs

The functions of the DEGs were determined by GO enrichment analysis. A total of 132 GO terms were enriched with DEGs identified between the oeCD36 and shCD36 groups, and each term was enriched with 2 genes. Among these terms, 91 biological process (BP), 26 cellular component (CC) and 15 molecular function (MF) terms were significantly enriched with DEGs (P<0.05) (Fig. 7A; Table EV2). Next, a KEGG pathway enrichment analysis was performed, and a total of 17 pathways were significantly enriched with DEGs identified between the oeCD36 and shCD36 groups (P<0.05). Most of the enriched pathways were metabolic pathways, FA metabolism pathways, amino acid metabolism pathways, endocytosis and other pathways (Fig. 7B; Table EV3).

**Figure 7.**
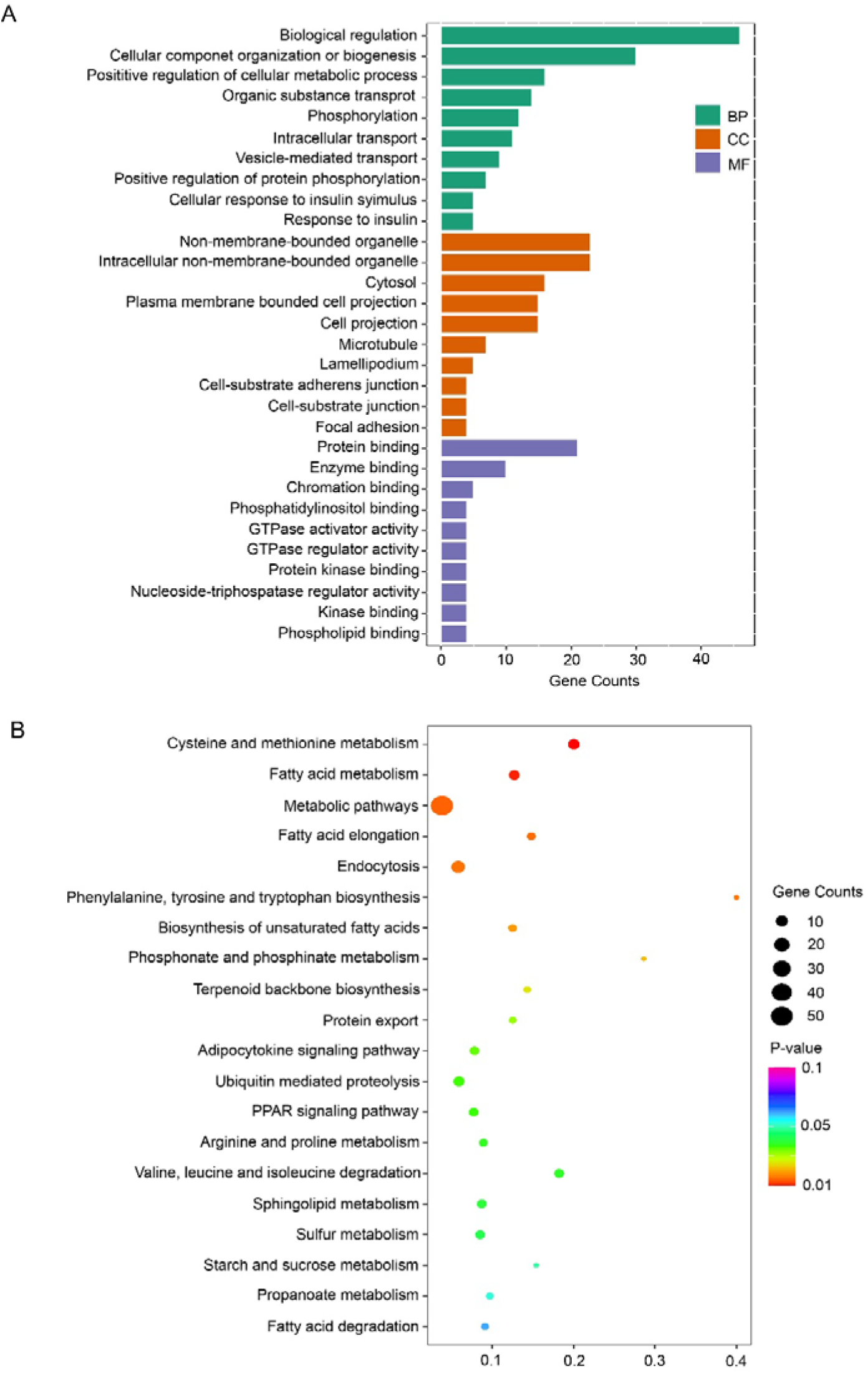
Gene Ontology (GO) and Kyoto Encyclopedia of Genes and Genomes (KEGG) enrichment analysis of the differentially expressed genes (DEGs). A. GO enrichment analysis of the DEGs. The top 10 biological process (BP), cell component (CC), and molecular function (MF) terms with DEGs identified between the oeCD36 and shCD36 groups. B. KEGG enrichment analysis of DEGs. KEGG pathways enriched with DEGs identified between the oeCD36 and shCD36 groups (Top 20).

### PPI network analysis of the DEGs between the oeCD36 and shCD36 groups

The results indicated that upregulated CD36 promoted the expression of *RXRA* through a positive feedback loop and inhibited the expression of *SLC38A1*, *GRB2* and *NCOA3*. The inhibition of *NCOA3* expression led to decreased expression of *WDFY3*, *PEAK1*, *PRRC2C*, and *UBR1*. Reduced expression of *GRB2* inhibited the expression of *WDFY3*, *PEAK1*, *NAV3* and *SEMA6A*. The decreased expression of *ASH1 L* further decreased the expression of *NCOA3*, *PRRC2C*, *UBR1*, *WDFY3*, *LYST*, *PEAK1*, *SLC4A7*, *MAP1B* and *ZZEF1*. Therefore, the differential expression of *CD36* affected the expression of the *NCOA3*, *ASH1 L*, and *PRRC2C* genes and thus affected the transport and degradation of lipids in CS2 cells (Figure 8).

**Figure 8.**
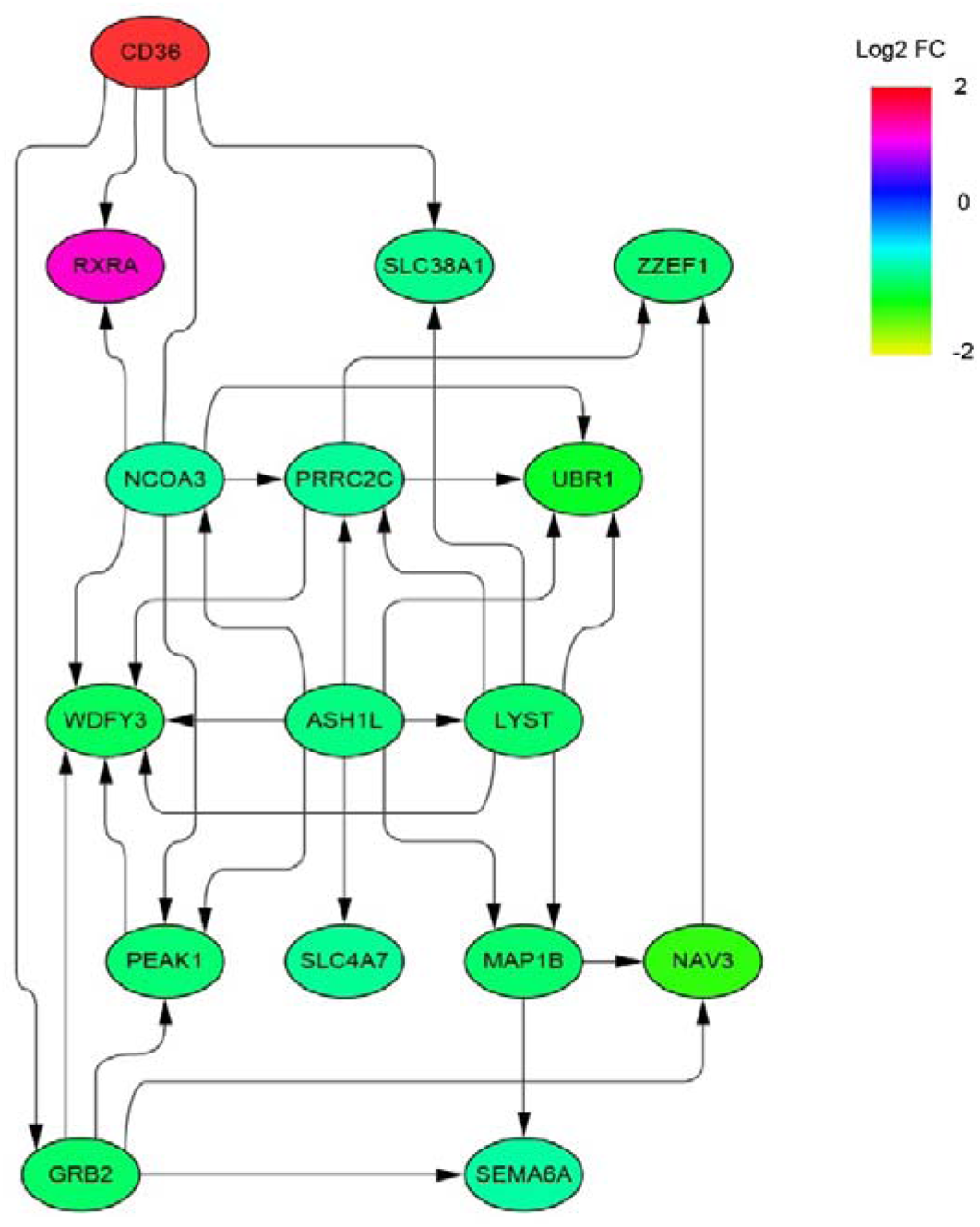
PPI-Network analysis of DEGs. A PPI network prediction analysis was performed to further explore the interaction between CD36 and DEGs. A network of DEGs identified between the oeCD36 and shCD36 groups was constructed with STRING and Cytoscape software.

## Discussion

Compared to other tissues, the skeletal muscle tissue of meat ducks consumes the most energy and is the centre of fat metabolism (Shaw *et al*, 2010). *RXRA* is one of the most widely distributed transcriptional regulators, and it participates in the expression of key genes in lipid metabolism (Kosters *et al*, 2013). Different RXR-selective agonists affected TGs in rat serum (Wan *et al*, 2000; Kliewer *et al*, 1992; Willy *et al*, 1995). The degree to which agonists cause acute hypertriglyceridaemia differs by the duration of agonist exposure (Peters *et al*, 1997). The accumulation of TGs in skeletal muscle is closely associated with insulin resistance and type 2 diabetes (Machann *et al*, 2004). This study found that *RXRA* overexpression led to a decrease in the content of TGs in CS2 cells, while interference with *RXRA* gene expression led to an increase in the content of TGs in CS2 cells. These results suggest that *RXRA* may regulate TG metabolism by binding to ligands that regulate TG transport in myoblast cells. This study also showed that *RXRA* promoted the FA β-oxidation of *PPARG* and reduced the accumulation of TGs in cells by increasing *PPARG* expression. *PPARG* can induce the expression of *ACSL 1*, which plays an important role in FA breakdown (Bervejillo *et al*, 2020). *ELOVL6* is a microsomal enzyme involved in the elongation of 12-, 14-, and 16-carbon FAs, a process regulated by *AMPK* signalling (Takashi *et al*, 2009; Sunaga *et al*, 2016). This study showed that *RXRA* overexpression promoted the gene and protein expression of *ELOVL6*, and interfering with *RXRA* expression inhibited the gene and protein expression of *ELOVL6* in CS2 cells. In addition, *RXRA* has been previously shown to play a key role in CHOL homeostasis, intestinal CHOL absorption, and bile acid synthesis (Willy *et al*, 1995). Rexone (LG268) activates RXRs (LXR and FXR) to form specific dimers, which inhibit cholesterol absorption and reverse cholesterol transport through the expression of the antiporter *ABC1* and the bile synthesis rate-limiting enzyme *CYP7A1* (Repa & J., 2000). Notably, *ABC1* is a target gene of RXRA (Selva *et al*, 2004; Nebel *et al*, 2009). Moreover, an analysis of an ABC1-knockout mouse model produced results similar to those of the RXR-knockout model. ACBA1 is a member of the ABC transporter family, and this member transfers CHOL from the periphery to ester-free apolipoprotein A1 to form high-density lipoprotein (HDL) (Brewer & Santamarina-Fojo, 2003). Overexpression of *ABCA1* in mice increased plasma HDL levels and prevented atherosclerosis (Singaraja *et al*, 2003). Therefore, *RXRA* plays an important role in regulating TG, CHOL and NEFA in CS2 cells, and these metabolites are closely related to lipid metabolism.

Bexarotene reduced dietary cholesterol absorption via *NPC1L1* and *CD13* expression in a study of hypertriglyceridaemic mice treated with bexarotene, which is an agonist of RXR (Lalloyer *et al*, 2006). This outcome was similar to the results of the present study. Therefore, we speculated that, in duck skeletal muscle cells, *RXRA* may accelerate the transport of TG and CHOL in myoblast cells through transporters such as *CD36*, *NPC1L1*, *CD13*, and *ABCA1*. Therefore, we predicted *RXRA* transcription factor-binding sites in the *CD36* promoter region. Our results showed that *RXRA* overexpression promoted the gene and protein expression of *CD36*; in contrast, interference with *RXRA* expression inhibited the gene and protein expression of *CD36* in CS2 cells. Moreover, we found that *RXRA* promoted the activity of the wt-CD36 promoter dual-luciferase reporter gene, as indicated with results obtain from an analysis of the dual-luciferase reporter gene expression relative to the mu-CD36 promoter dual-luciferase reporter gene expression (with the five predicted binding sites deleted). This finding also suggested that *RXRA* regulated the transcription of *CD36* by targeting the *CD36* promoter activity, and this increase in *CD36* gene expression led to the transport of TG, CHOL and NEFA in myoblasts. To confirm this speculation, *CD36* overexpression and interference vectors were constructed and used explore the effect of *CD36* expression on the TG, CHOL and NEFA contents in CS2 cells.

The surprising results showed that *CD36* had the same function as *RXRA* and reduced the contents of TGs, CHOL and NEFAs in CS2 cells. Previously, Ibrahimi established a model of *CD36* overexpression by activating the creatine kinase (MCK) promoter in mouse muscle, and the results showed that the transgenic mice lost weight and adipose tissue and exhibited significantly reduced TG, CHOL, FAs, and very-low-density lipoprotein (VLDL) levels (Ibrahimi *et al*, 1999). These results were similar to those of the present study, suggesting that *CD36* plays an important role in fat metabolism in myoblasts. Koonen transfected the *CD36* gene with adenovirus in mouse models of obesity and found that increasing the expression of *CD36* promoted the formation of intrahepatic cholesterol esters, promoted the uptake of FAs in the liver, and increased the TG level (Koonen *et al*, 2007). These results differed from those in our study, which may be due to the analyses of different tissues. Notably, long-chain FAs are essential for lipid metabolism (Tran *et al*, 2016). The results of our study showed that CD36 inhibited the accumulation of SFAs. The human body can synthesize SFAs, but some unsaturated FAs must be obtained from external sources, i.e., food. However, the increase in SFAs in animals can lead to cell inflammation, steatosis and insulin resistance (Liu *et al*, 2019). Moreover, endogenous biosynthesis of myristic acid is indeed far lower than the amounts consumed from dietary sources (Legrand & Rioux, 2010). Evidence suggests that myristic acid is not only beneficial to endothelial cells but is also crucial to the activation of platelet-reactive protein-1, and its level can be regulated by *CD36* uptake from medium (Isenberg *et al*, 2007). High-MUFA meals have beneficial effects through postprandial lipidaemia responses, and cardiovascular risk factors were reduced and lipid profiles improved after MUFA intervention (Lopes *et al*, 2016). Oleic acid is the most abundant MUFA in humans (Oi-Kano *et al*, 2007). When we knocked down the expression of the *CD36* gene in CS2 cells, MUFA and oleic acid accumulation was increased, while the overexpression of the *CD36* gene decreased the content of oleic acid but had no effect on MUFA levels. The increase of MUPAs in duck meat is a favourable reference for human dietary FA composition. Finally, we found that *CD36* expression exerted no effect on PUFA content, but after inhibiting *CD36* expression, the contents of eicosapentaenoic acid (EPA) in the CS2 cells increased. Dietary PUFAs play a protective role in improving cardiovascular disease by controlling the synthesis and oxidation of SFAs and MUFAs (Jump, 2011). The EPA in fish oil is thought to be important for inducing hypotriglyceridaemic effects in humans (Ebbesson *et al*, 2007). We speculated that other genes or pathways regulate the content of long-chain FAs in cells and that the mechanism by which changes in *CD36* expression lead to a reduction of long-chain FA contents requires further study.

Many protease receptors and transporters are involved in lipid metabolism. These receptors and transporters are regulated by the signal transduction pathway, forming a complex and fine-tuned regulatory network that maintains the lipid metabolism balance in cells and the whole organism. In this study, CS2 cells were used as the research objects to analyse the correlation between *CD36* overexpression and *CD36* interference, with 71 DEG candidates identified. These DEGs were enriched in the 132 GO terms, and we found that GO terms related to organic matter transport and protein phosphorylation were enriched with DEGs. Interestingly, *NCOA3*, *GRB2*, *SLC38A1* and *SLC4A7* are key genes involved in vehicle-mediated transport, and *RXRA*, *PEAK1*, *MAP1B* and *SEMA6A* are key genes involved in the response to insulin. Differentially expressed genes (DEGs) were significantly enriched in 17 pathways, among which GRB2 was engaged in FA elongation, endocytosis and PPAR signalling pathways, while NCOA3 was a key gene in the insulin signalling pathway. GRB2 is a widely studied cell signalling adaptor protein with particular importance in the *PI3K/AKT* pathway (Zhi *et al*, 2008). Inhibition of *GRB2* expression attenuated fat accumulation, oxidative stress and apoptosis by regulating insulin signalling (Shan *et al*, 2013, 2). The PPI network interaction analysis performed in this study revealed that increasing the expression of *CD36* inhibited the expression of the *GRB2* gene; therefore, *CD36* may reduce the lipid accumulation in CS2 cells by regulating the expression of the *GRB2* gene and promoting the excretion of intracellular organic lipids. Surprisingly, the expression of *NCOA3* was also downregulated. *NCOA3* constitutes one-third of the steroid receptor coactivator (SRC) family in the body, enhancing nuclear transcriptional capacity, including oestrogen receptor (ER) transcription. *NCOA3* plays an important role in fat accumulation, and inhibiting the expression of *NCOA3* can reduce the TG content and inhibit lipid accumulation in porcine adipocytes (Haiyin *et al*, 2017). *MAP1B* is the primary phosphorylated substrate of the serine/threonine kinase glycogen synthase kinase-3β (GSK-3β) in differentiated neurons, and GSK-3β plays an important role in the insulin resistance pathway (Barnat *et al*, 2016). *RXRA* regulates the expression of *CD36*, and the DEGs identified in this study that increased the expression of *CD36* also promoted the increased expression of *RXRA*, which may have been due to the inhibition of *NCOA3* expression, which promoted *RXRA* expression through a negative regulatory feedback mechanism. When *CD36* was overexpressed, the expression of *MOB2*, *SAA2* and *PKIA* was increased, but that of *SLC38A1* and *SLC4A7* was reduced. *MOB2*, a Mps one binder (MOB) family protein, regulates NDR/LAT kinase family members and exerts various biological functions, such as subcellular localization, protein–protein interaction, and cell migration (Hergovich, 2011; Jiang *et al*, 2020). *SAA2* is a member of the apolipoprotein family and plays a vital role in CHOL clearance (Manley *et al*, 2006). *PKIA* is a member of the cAMP-dependent protein kinase (PKA) inhibitor family, which has been shown to inhibit the activity of a PKA subunit (T Güttler *et al*, 2010, 1). cAMP has also been shown to regulate exocytosis in various secretory cells, with PKA producing the main signal for this regulatory process (Seino & S., 2005). These findings comprise favourable evidence for lipid efflux from CS2 cells in this study. The expression of *SLC38A1*, an important transporter of glutamine and a precursor of the neuronal synaptic transmitter glutamate, was significantly upregulated in the brown adipose tissue of mice (Horie *et al*, 2017). *SLC4A7* is an important factor in maintaining cellular homeostasis during the macrophage phagocytosis (Sedlyarov *et al*, 2018). Therefore, we speculated that when the CS2 cell membrane composition is changed, *SLC4A7* regulates the homeostasis of intracellular pH to maintain cellular homeostasis. These results indicate that *CD36* increases lipid efflux from CS2 cells by increasing the expression of *MOB2*, *SAA2* and *PIKA* and that *SLC4A7* is a critical factor in maintaining the stability of the cell composition.

Considering the results of our study, we propose a model showing the inhibition of lipid accumulation in CS2 cells as mediated by *RXRA* and *CD36*. In this model, *RXRA* not only independently regulates lipid beta-oxidation but also precisely regulates *CD36* expression to inhibit lipid accumulation in CS2 cells. Therefore, both *RXRA* and *CD36* contribute to lipid reduction in CS2 cells. Although we have established that *RXRA* and *CD36* comprise a regulatory network, other components in CS2 cells may be involved in this network. *CD36* regulates CS2 cell lipid metabolism by regulating endocytosis, FA synthesis and amino acid metabolism. Moreover, cell homeostasis is maintained through the regulated expression of intracellular cytokines. *GRB2*, *MAP1B*, *SLC38A1*, *SLC4A7*, *NCOA3*, *PKIA*, *MOB2*, *SAA2* and *RXRA* are involved in the regulation of CS2 cell lipid outflow induced by *CD36*. In summary, we found that *RXRA* reduced lipid accumulation in CS2 cells and regulated the accelerated outflow of FAs mediated through *CD36*, providing a molecular framework that demonstrates that *RXRA* regulates lipid accumulation.

## Materials and Methods

### Ethics approval

The animal experiment was reviewed and approved by the Institutional Animal Care and Use Committee of Anhui Agriculture University (no. SYDW-P20190600601). The experiments were performed in accordance with the Regulations for the Administration of Affairs Concerning Experimental Animals and the Standards for the Administration of Experimental Practices. Tissues were collected from 13-day-old duck embryos. The duck embryos were taken from a farm in Qiangying, Anhui.

### Primary myoblast separation and culture

The separation and culture methods used to obtain myoblasts (CS2 cells) were described in a previous study (Shan *et al*, 2014). Dulbecco’s modified Eagle’s medium/nutrient mixture F-12 (DMEM/F12, VivaCell, 01-172-1ACS), containing 10% foetal bovine serum (FBS, VivaCell, C04001-500) and a 100 U/mL penicillin– streptomycin solution (VivaCell, 03-0321-1B), was used to culture CS2 cells.

### Vector construction

RNA from duck myoblasts was extracted using the TRIzol (Invitrogen, 15596018) method. RNA was reverse transcribed into cDNA according to the instructions in a first-strand synthesis premix kit (Vazyme, R122-01). Full-length *RXRA* (XM_027471073.2) was amplified with specific primers, including the NheI (NEB, R3131S) and HindIII (NEB, R3104 V) restriction sites. The pBI-CMV3 vector and an *RXRA* fragment were digested with NheI and HindIII. OeRXRA was constructed by linking the purified pBI-CMV3 vector with the RXRA fragment. OeRXRA was transfected into CS2 cells using ExFect transfection reagent (Vazyme, T101-01). The transfection conditions were ExFect:oeRXRA= 2:1. Four interference targets were identified based on the full-length *RXRA* sequence, and upper and lower primers were designed, including the AgeI (NEB, R3552S) and EcoRi (NEB, R310T) restriction sites at the 5’ and 3’ ends, respectively. The primers were annealed to form double-stranded oligomers and attached to the PTSB-SH-copGFP-2A-PURO skeleton to construct short interfering RNAs: shRXRA1, shRXRA2, shRXRA3, and shRXRA4. These four interference vectors against RXRA were transfected into CS2 cells. After 24 h, cell samples were collected, and the RNA was extracted and reverse transcribed into cDNA. The effects of each interference vector on *RXRA* expression were quantitatively determined by RT–PCR (Vazyme, Q711-02). The vector with the highest interference effect was named shRXRA and used for all subsequent experiments. The above methods were also used to construct *CD36* (XM_038183702.1) overexpression and interference vectors. The primer synthesis in this study was completed by Shanghai Sangon Biotechnology Co., Ltd. The primers for preparing the overexpression and interference vectors of the *RXRA* and *CD36* genes can be found in the Supplementary data (including Table EV4 and Table EV5). A wild-type CD36 promoter luciferase reporter vector (wt-CD36) and binding site mutant vector (mu-CD36) were synthesized by Shanghai Gene Pharma Co., Ltd. and cloned into a GPL3-Basic vector to form wt-CD36 and mu-CD36. A PGL3-RL vector was purchased from Shanghai Gene Pharma Co., Ltd.

### Luciferase reporter assay

The mature 2000-bp sequence of the *CD36* promoter was selected for transcription factor prediction using PROMO (http://alggen.lsi.upc.es/cgi-bin/promo_v3/promo/promoinit.cgi?dirDB=TF_8.3) software. PROMO predicted five targeted binding sites for *RXRA* and the *CD36* promoter region. The complete *CD36* promoter sequence (wt-CD36) and the *CD36* promoter sequence with five target sequences missing (mu-CD36) were submitted to Gemma Biosynthesis, which ligated them into a GPL3-Basic cloning vector (the complete promoter sequence can be viewed in the Supplementary file). OeRXRA/oeNC/wt-CD36/mu-CD36/GPL3-Basic was cotransfected into CS2 cells, and a Renilla luciferase vector (PGL-RL) was used to correct the transfection efficiency data. The firefly (hluc+) and Renilla (hRluc) luciferase intensities were detected with a Dual-Glo luciferase assay system (Promega, E1960) according to the manufacturer’s protocols; hluc+ luciferase was used as the reference to correct for variations in transfection efficiency, and the relative luciferase activity was calculated with the following equation: relative luciferase activity=hRluc/hluc+. A Bio-Tex microplate reader was used to detect the fluorescence values and determine whether the promoter of the *CD36* gene contained binding sites for RXRA.

### Lipid metabolism-related gene expression in CS2 cells

After 24 h of transfection, the CS2 cells that had been transfected with vectors were collected, and TRIzol was used to lyse the CS2 cells before total RNA extraction. cDNA was reverse transcribed from total RNA using a cDNA synthesis kit. RT–PCR (ABI7500, Thermo, America) was performed with qPCR mix. With GAPDH as the reference gene, RT–PCR was used to detect the mRNA levels. We used the 2^-ΔΔct^ method to calculate the relative expression of the genes expressed in CS2 cells. Primers for RT–PCR can be found in Table EV6.

### western blotting

Cells transfected with plasmids were harvested 24 h after transfection. The cells were then resuspended in RIPA buffer with protease inhibitors (Meilunbio, MA0151). The cell lysate was removed by centrifugation at 12000 rpm/min for 10 min at 4°C, and the supernatant was collected. The BCA (Vazyme, E112-02) protein quantitative analysis method was used to determine the total protein concentration in the samples. For western blotting (Bio–Rad, PowerPac™ HV), the same amount of protein was separated by SDS–PAGE and then transferred to PVDF (Thermo-Fisher, 88518) membranes with a gel-to-membrane transfer system. After blocking with 1×PBS buffer (VivaCell, C3580-0500) containing 1% bovine serum albumin (BSA) (Servicebio, GC305006) for 2 h, the PVDF membrane was washed with a TBST (Sangon Biotech, B040126) solution. Then, the membrane was incubated with primary antibody (diluted in 1% BSA) at 4°C for 12 h. Primary antibodies against RXRA (YN0018), GAPDH (YM3215), FABP4 (YM0013), ELOVL6 (YT1540), and ACSL1 (YN0827) were all purchased from ImmunoWay, China; PPARG (16643-1-AP) was purchased from Proteintech, China; and CD36 (bs-8873R) was purchased from Bioss. The membrane was incubated with a secondary antibody (ImmunoWay) (1:1000 diluted in 1% BSA) for 2 h. The signal was visualized using an enhanced chemiluminescence kit (Vazyme, E412-01). The relative intensities of the bands were quantified with ImageJ software.

### Analyses of the intracellular TG, CHOL, and nonesterified fatty acid (NEFA) content

Following the respective manufacturer’s instructions, we used kits to determine the TG (Applygen, E1013-105), CHOL (Applygen, E1015-105), and NEFA (Nanjing Jiancheng, A042-2-1) contents. The CS2 cells were collected 24 h after transfection with the overexpression plasmid or the interference plasmid. Different samples were adjusted to the same cell level through BCA protein quantification, and the test for each model was repeated three times. The TG, CHOL, and NEFA levels in the cells were adjusted according to the protein content. A microplate reader was used to determine the concentrations of TGs, CHOL, and NEFAs.

### Fatty acid (FA) targeting analysis

Fatty acids (FAs) were extracted from CS2 cells 24 h after transfection. The cells were trypsinized after washing 3 times with PBS. Cell pellets were collected through centrifugation. The cells were placed in a 2-mL centrifuge tube, 1 mL of chloroform:methanol (2:1) solution and 100-mg glass beads were added, and then, the samples in the centrifuge tube were homogenized with a tissue grinder, shaken at 60 Hz for 1 min, and centrifuged at 12000 rpm at 4°C for 5 min. All the supernatant was placed in a 15-mL centrifuge tube, and 2 mL of a 1% sulfuric acid methanol solution was added, mixed well and shaken for 30 s. The sample was allowed to stand for 5 min, and then, 5 mL of H_2_O was added to wash the sample, which was then centrifuged at 3500 rpm at 4°C for 10 min. Next, 700 μL of the supernatant was added by pipetting into a new 2-mL centrifuge tube, and 100 mg of anhydrous sodium sulfate powder was added to remove excess water from the sample, which was shaken and mixed for 30 seconds before centrifugation at 12000 rpm for 5 min. Then, 300 μL of the supernatant was pipetted into a new 2-mL centrifuge tube, and 15 μL of 500 ppm methyl salicylate was added. An ester was used as the internal standard, and after the standard was added, the sample was shaken and mixed for 10 s. Finally, 200 μL of the supernatant was added to a test bottle. The operating conditions were as follows: For chromatography, a Thermo TG-FAME capillary column (50 m*0.25 mm and an internal diameter of 0.20 μm) with an injection volume of 1 μL at a split ratio of 8:1 was used. The inlet temperature was 250 ℃, the ion source temperature was 300 ℃, and the transmission line temperature was 280 ℃. The starting programme temperature was 80 ℃, which was maintained for 1 min; then, the temperature was increased to 160℃ at a rate of 20℃/min, and it was maintained for 1.5 min; then, it was increased to 196 ℃ at 3 ℃/min and maintained for 8.5 min; and finally, the temperature was increased to 250 ℃ at 20 ℃/min and maintained for 3 min. The carrier gas was helium, and the carrier gas flow rate was 0.63 mL/min. An electron impact ionization (EI) source, SIM scanning mode, and electron energy of 70 eV were the mass spectrometry conditions for this experiment. The column was cleaned after 4 samples were run. The FA contents were calculated through peak area normalization. The 38 standard FAs were measured, and then, the number of times that the 38 FAs were detected was determined. The concentration and the added volume of the internal reference FA were known, the content of each tested FA was calculated according to the quantity of the internal reference FA, and the proportion of the tested FA to the total amount of FA was determined. The experiment was repeated 3 times, providing 3 biological replicates, and each replicate was measured 3 times.

### RNA extraction and RNA sequencing (RNA-seq)

The CS2 cells were collected 24 h after transfection with the oeCD36 or shCD36 group plasmids. Total RNA was extracted from 6 cell samples using TRIzol reagent according to the manufacturer’s instructions. RNA sequencing (RNA-seq) was performed by Shanghai Shenggong Biotechnology Co., Ltd. (Shanghai, China).

### Statistical analysis

The experimental data are shown as the means ± standard error of the mean. Significant differences were identified with a P value < 0.05. GraphPad Prism 9 software (GraphPad Software Inc., La Jolla, CA) with a 2-tailed (unpaired) t test was used for the data analysis. The RNA-seq data were aligned to the reference genome using HISAT2, and gene expression was assessed using StringTie and known gene models. Differentially expressed genes (DEGs) were identified using the DESeq R package functions estimate SizeFactors and nbinomTest. A Q value < 0.05 and log2-fold change (FC) >1 or log2 FC < -1 were set as the thresholds for determining significant differential expression. Based on the gene expression levels in the different samples, principal component analysis (PCA) diagrams were generated with ggplot2. Volcano plots showing DEGs between different groups were drawn with ggplot2. Gene Ontology (GO) enrichment analysis was performed using topGO, and Kyoto Encyclopedia of Genes and Genomes (KEGG) pathway enrichment analysis was performed by cluster profiling. The significance level for the enriched GO terms and KEGG pathways was set to P<0.05. Protein–protein interaction (PPI) networks were constructed using the STRING online database based on DEGs and visualized using Cytoscape software.

## Acknowledgements

This work financially supported by Major Scientific and Technological Special Project in Anhui province (201903a06020018). We thank Qiangying Duck Industry for providing duck embryo eggs, Zhihui Zhao for providing the pBI-CMV3 vector.

## Credit authorship contribution statement

Pan Ziyi: Conceptualization, Methodology, Software, Writing - Review & Editing. Li Guoyu: Methodology, Formal analysis. Du Guoqing: Methodology, Validation. Wu Dongsheng: Software, Validation. Li Xuewen: Formal analysis, Investigation. Yu Wang: Data Curation, Writing - Original Draft. Junxian Zhao: Data Curation, Writing - Original Draft. Xiran Zhang: Data Curation, Writing - Original Draft. Chen Xingyong: Visualization, Conceptualization. Zhang Chen: Conceptualization, Supervision. Jin Sihua: Conceptualization, Supervision. Geng Zhaoyu: Conceptualization, Writing - Review & Editing, Project administration, Funding acquisition.

## Data availability

Data and code related to this paper may be requested from the authors or supplied in the manuscript.

